# Inverted transflection spectroscopy of live cells using metallic grating on elevated nanopillars

**DOI:** 10.1101/2023.09.19.558443

**Authors:** Aditya Mahalanabish, Steven H. Huang, Gennady Shvets

## Abstract

Water absorption of mid-infrared (MIR) radiation severely limits the options for vibrational spectroscopy of the analytes – including live biological cells – that must be probed in aqueous environments. While internal reflection elements, such as attenuated total reflection prisms and metasurfaces, partially overcome this limitation, such devices have their own limitations: high cost, incompatibility with standard cell culture workflows, limited spectral range, and small penetration depth into the analyte. In this work, we introduce an alternative live cell biosensing platform based on metallic nanogratings fabricated atop elevated dielectric pillars. For the MIR wavelengths that are significantly longer than the grating period, reflection-based spectroscopy enables broadband sensing of the analytes inside the trenches separating the dielectric pillars. Because the depth of the analyte twice-traversed by the MIR light excludes the highly absorbing thick water layer above the grating, we refer to the technique as Inverted Transflection Spectroscopy (ITS). We demonstrate the analytic power of ITS by measuring protein concentrations in solution. The ability of ITS to interrogate live cells that naturally wrap themselves around the grating is also exploited to characterize their adhesion kinetics.

---

All-optical label-free sensors have revolutionized the field of analytic life sciences, enabling the detection and quantification of biological objects such as proteins, DNA, and live cells. These biosensors have led to significant advances in drug discovery and clinical diagnostics. Mid-infrared (MIR) spectroscopy has emerged as a popular tool for biosensing because of its unique ability to resolve characteristic vibrational fingerprints of the constituent biological molecules. It has already gained significant attention due to its non-invasive, label-free, and rapid detection capabilities. For example, Fourier-Transform Infrared (FTIR) spectroscopy has been used in the past to characterize proteins and polypeptides.^1–4^ However, MIR spectroscopic measurements suffer from several important limitations that restrict their usefulness for analyzing biological material in aqueous solution, e.g., in patient-derived biofluids.^5,6^

The leading issue is the strong attenuation of mid-IR light in water, which limits the sensitivity of the spectroscopic technique. Because of the water-specific absorption due to OH bending vibrational band around *ω*_H–O–H_∼ 1,650 cm^−1^, this issue is particularly severe in the biologically important spectral range 1,600 cm^−1^ < *ω* < 1,700 cm^−1^ associated with Amide I protein vibrations. As the result, the optical path length needs to be short (typically, smaller than 6-8 μm) and the analyte concentration needs to be high (typically, larger than 3 mg/mL for proteins^3^) to ensure reliable transmission-based measurements. Water absorption similarly affects another popular approach to collecting the spectra in reflection: the so-called transflection-mode spectroscopy ^7–9^, where MIR light passes through the analyte and reflects off a substrate coated with infrared-reflective layer. Longer optical paths and higher sensitivity can be achieved using brighter MIR light sources, such as quantum cascade lasers (QCL), albeit at the expense of the increased complexity of the setup required to maintain its constant temperature^10–12^. Using a high-index prism to measure attenuated total reflection (ATR) of MIR light circumvents water absorption^1,13–15^ but limits the penetration depth into analytes to a fraction of a micron,^15^ thus reducing the sensitivity of the approach.

In addition to characterization of liquid biological samples, MIR spectroscopy has shown great potential in cellular studies. MIR spectroscopy-based cytology based on the quantification of vibrational fingerprints of cellular molecular constituents (e.g., proteins, lipids, carbohydrates, and nucleic acids) provides an excellent tool for rapid differentiation between different cell types^16–18^, as well as the detection of cellular responses (e.g., apoptosis under cytotoxic conditions) under various influences (e.g., environmental changes, drugs, and other stimuli^19^). For FTIR spectroscopy of live cells, the use of attenuated total reflection – FTIR (ATR-FTIR),^13,15,20–23^ and transmission measurements using thin microfluidic devices^24,25^ have been reported. Use of shallow transmission cells can impose mechanical stress on cells, while the ATR-FTIR approach suffers from a very shallow penetration depth into a cell^23^. From a practical standpoint, neither of the two approaches can be readily integrated with standard cell culture workflows, making the transition to high-throughput cell assaying based on MIR spectroscopy challenging.

Recently, our group has developed metasurface-enhanced infrared spectroscopy (MEIRS) and used it in a variety of life science applications, including sensing single protein layers,^4^ distinguishing between and capturing of different cell types,^26,27^ and, more recently, measuring the MIR spectra of live cells in real time.^28–30^ Metasurfaces, made up of arrays of metallic nanoantennas, rely on plasmonic resonances to enhance and localize optical field around the nanophotonic structures. The coupling of molecular absorption to the plasmonic resonances enables characteristic molecular vibrations to be visible in the reflected light. By integrating metasurfaces with a traditional FTIR system, our group demonstrated the use of MEIRS as a cellular assay technique to study cell adhesion and detachment, responses of live cells to chemotherapeutics,^30^ as well as cholesterol depletion from cellular membrane.^28^ One drawback of any resonant metasurface is its spectral selectivity: the relatively narrow bandwidth of the resonance can be insufficient for covering the entire fingerprint spectral range. Although multi-resonant metasurfaces with broad spectral coverage have also been used for biosensing applications,^31^ such multi-resonant metasurfaces rely on nanoantennas of different geometries supporting different resonance modes and having different nearfield distributions. As a result, signals from different wave-numbers originate from different regions of the metasurface that have limited spatial overlap. This is not a problem for the analytes distributed uniformly across the metasurface. However, because cells have different organelles, cellular samples are highly heterogeneous.

Therefore, utilizing different sensing volumes for interrogating different vibrational bands (e.g., lipids, proteins, carbohydrates) can confound the interpretation.

In this work, we propose an alternative concept for a broadband optical device based on non-resonant plasmonic nanostructures (see Fig.1(a)) that enables measuring MIR spectra of cells and biomolecules in aqueous solution. Unlike the MEIRS approach, which is based on resonant metasurfaces, the approach described below does not rely on the localized resonance of plasmonic nanostructures and thus has a much broader spectral sensing range. We refer to the proposed technique as Inverted Transflection Spectroscopy (ITS) because of its similarity to traditional transflection-mode^7–9^ measurements, as explained below.

**Figure 1:**
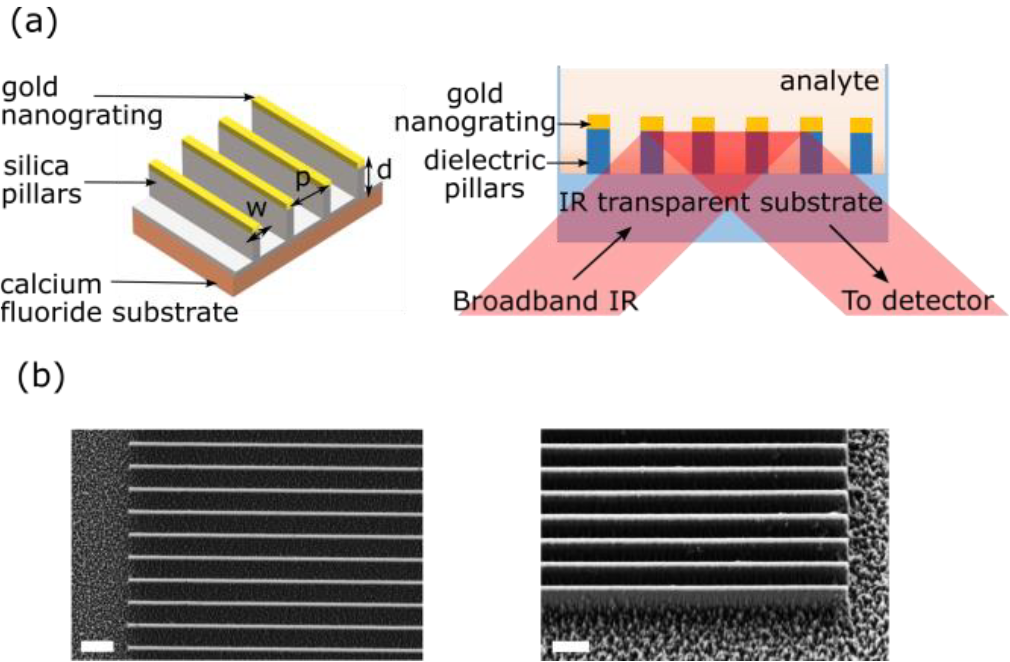
(a) Schematic drawing of the Inverted Transflection Spectroscopy (ITS) device: MIR light polarized along the grating direction and incident from the substrate side reflects from the grating, passes twice through shaded region. (b) SEM images of a fabricated sample. Parameters of the elevated nanograting: *w* = 200 *nm, p* = 1. 35 *μm, d* = 830 *nm*. (Left) top view, scale bar: 2 μm; (Right) the same sample viewed at 35° angle. Scale bar: 1 μm.

## Theoretical Background

The ITS measurement is based on the device schematically shown in Figure 1(a) comprising a periodic array of gold strips atop silica pillars grown on an IR transparent substrate. The device functions similarly to a wire grid polarizer under normal incidence, where light with electric field polarized parallel to the gratings is reflected (with minimal transmission) when the grating periodicity *p* is much smaller than the incident wavelength of light in vacuum *λ* ≡ 2*πc*/*ω*. Analytic expressions for the reflection coefficients 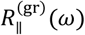 and 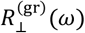 (with incident light polarized parallel and perpendicular to the grating direction, respectively) from a conducting plane with slits are given as follows^32^

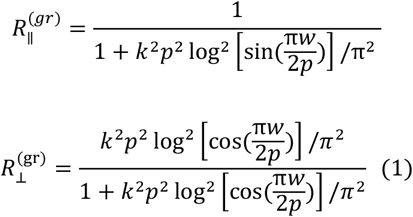

where *k* = 2*πn*_eff_/*λ* is the wavenumber of the incident wave in the embedding medium with a refractive index *n*_eff_ and *w* is the width of each conducting strip. Geometric parameters of the elevated nanograting are defined in the left panel of Fig.1(a).

It follows from Eq.(1) that in the long-wavelength limit (defined as *k p* ≪ 1), 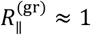 and 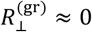. The regions of the trenches separating the nanopillars with the height *d* are twice-traversed by the ∥-polarized component of the incident light. Therefore, the total measured reflectance *R*_∥_(*ω*) accounts for the light absorption inside the analyte layer whose effective volume is controlled by the geometry of the elevated nanograting.

In contrast to conventional transflection-mode measurements, where the incident light beam traverses the entire analyte (i.e., incident from the top in Fig.1(a)), the measurement mode considered here correspond to the light beam incident from the inverted direction: hence the designation of Inverted Transflection Spectroscopy (ITS). On the other hand, the ⊥-polarized component of the incident beam propagates past the elevated nanograting without reflections and is subsequently absorbed by water.

Therefore, by limiting the sampled region to a small but finite depth *d*, the inverted geometry enables a transflection-mode measurement that would be impossible to carry out from the opposite direction because of strong water absorption of MIR light.

Crucially, any analyte contained within the trenches – up to the distance *d* above the substrate – can be analyzed through ITS by measuring the reflectance *R*_∥_(*ω*), which is a function of 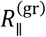 and absorbance of the analyte. Consequently, the total measured reflectance *R*_∥_(*ω*) depends on (a) the properties of the analyte, and (b) the optical path length *l* ∼ 2*d* through the sampled layer (which includes the silica pillars and the analyte) controlled by the height *d* of the silica pillars. In particular, attenuation of the MIR light in water can be limited to a few micrometers by keeping the pillar height small, making the proposed structure a useful device for measuring aqueous samples, as well as live cells in culture medium.

An important factor limiting the available spectral range of the present implementation of the ITS device to *ω* > 1,400 cm^−1^ is the presence of silica lattice vibrations modes^33^ in the 1,000 cm^−1^ < *ω* < 1,350 cm^−1^range grey-shaded in Figure 2(d). These reflectance dips are caused by the silica dielectric pillars underneath the gold gratings, as well as a thin layer of silica (∼200 nm) on the CaF_2_ substrate. The reduced raw (un-normalized) transflectance signal in the silica absorption band makes it essentially unusable for quantifying analyte-related spectral features. Therefore, in the rest of the article this spectral region is excluded from all spectra.

**Figure 2:**
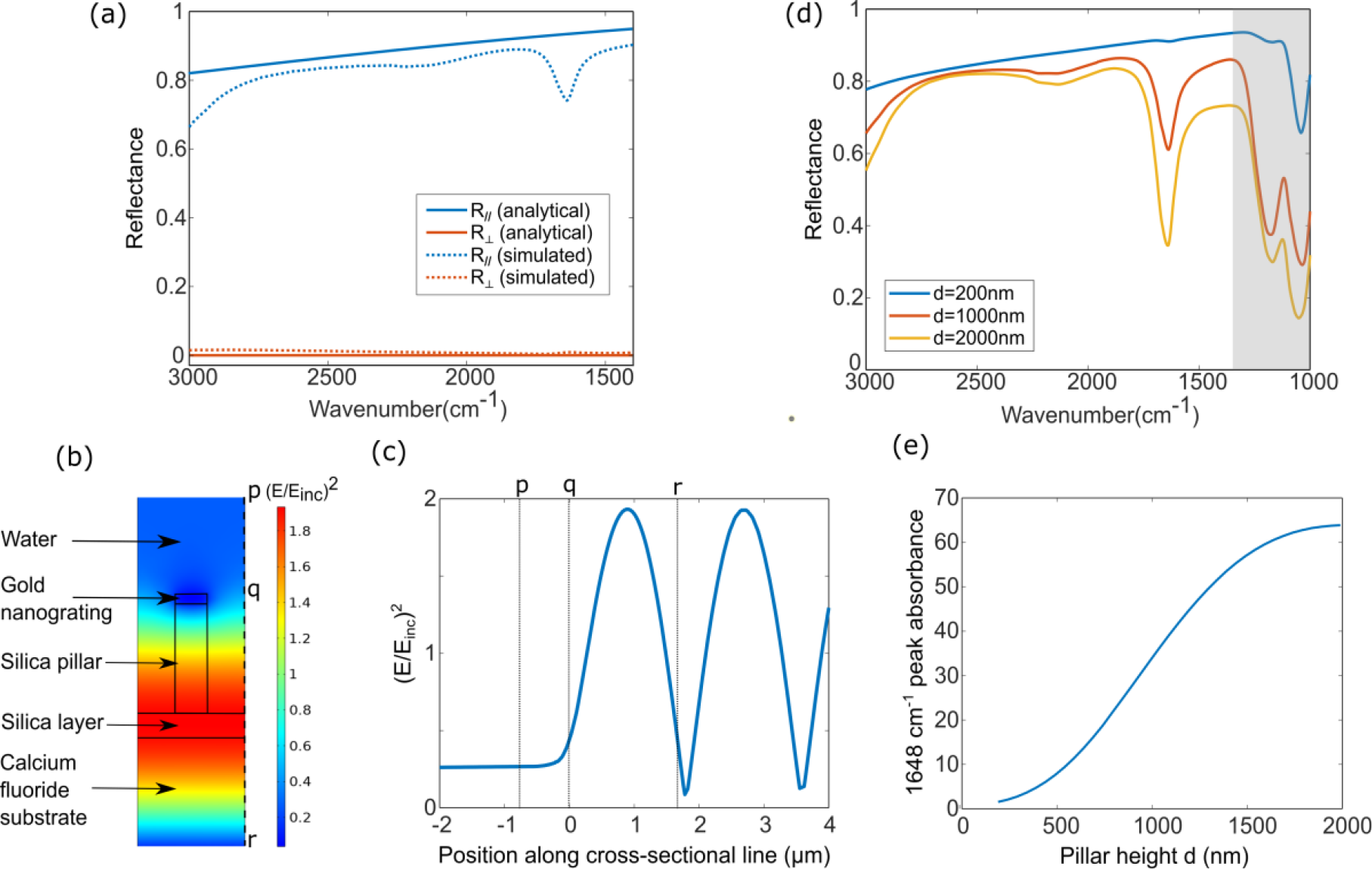
Simulations of ITS for pure water as analyte. (a) The analytic grating reflectance 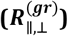 and simulated total (*R*_∥,⊥_) reflectance from the water-filled elevated nanogratings device. Light is normally incident through the CaF_2_ substrate with two polarizations: ∥ or ⊥ to the grating. (b) Normalized field intensity for ∥-polarized light with *λ* = 5 μm. (c) Field intensity along the dotted line marked in (b). Points of references p, q and r are marked in (b) and (c). (d) ITS spectra for 3 pillar heights: d = 200 nm, 1000 nm, and 2000 nm; with the greyed-out region marking the silica absorbance bands (e) Variation of water absorbance at *ω*_H–O–H_ ≈ 1, 650 *cm*^−1^, as seen from the ITS spectra, for pillar heights 200 nm < *d* < 2, 000 nm. Grating period: *p* = 0. 675 μm, height *d* = 830 nm and width *w* = 200 *nm*.

*Example*: *Water-filled ITS device*. To illustrate the working of the ITS approach, we begin by examining the simplest case of pure water as an analyte. Analytic calculations (see Eq.(1)) of 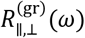 and numerical simulations (using COMSOL Multiphysics commercial software package) of 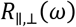 were carried out for the following parameters of the elevated grating: *p* = 675 nm, *w* = 200 nm, and the grating height *d* =830 nm. For the analytic calculations (solid lines in Fig.2(a)) we have chosen *n*_eff_ = 1.3as the frequency-independent average refractive index of water and silica. On the other hand, frequency-dependent tabulated refractive indexes of water and silica are used for the COMSOL simulations exhibited as dashed lines in Fig.2(a). The considered spectral range 1,350 *cm*^−1^ < *ω* < 3,000 *cm*^−1^ contains several important biological MIR fingerprints. Specifically, the CH_2_/CH_3_ stretching modes in the 2,800 cm^-1^ – 3,000 cm^-1^ region are attributed to lipids^34^, amide features in the 1,500 cm^-1^ – 1,700 cm^-1^ region are attributed to a combination of C=O stretching, N-H bending and C-N stretching in the amide backbone of proteins^35^, and several smaller features are attributed to phosphates (∼ 1,240 cm^−1^ : not examined here) and carbohydrates (1,000 – 1,500 cm^-1^).^36^ For the ⊥-polarized incident light, we observe that *R*_⊥_(*ω*) ≈ 0 for all frequencies, as expected because 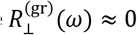. For the ∥-polarized light, 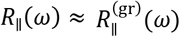 for most frequencies, except for the spectral region in vicinity of *ω* ∼ *ω*_H‣O–H_, where water is strongly absorbing. Deviations of *R*_∥_ from 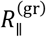 is attributed to trans-flection-mode absorbance *A*(*ω*): the ∥-polarized MIR light travels through the silica pillar/analyte layer, reflects from gold, and travels back through the silica pillar/analyte layer for a second time, before being collected at the detector.

Interpreting the resulting transflection spectra *R*_∥_(*ω*) requires considering several effects specific to ITS. First, because the analyte is in the form of a thin film (*d* < *λ*), there is an electric field standing wave (EFSW)^37–39^ formed within the analyte layer. This EFSW effect causes the absorbance to change non-linearly with respect to the sample thickness. Therefore, the Beer-Lambert law, which predicts a linear relationship between absorbance and the analyte depth *d*, no longer holds because the grating structure produces a complex wave intensity pattern inside the analyte/silica pillar layer. The resultant intensity distribution produced by the interference between the incident and reflected waves from the grating is shown in figures 2(b,c), with figure 2(c) showing the electric field intensity variation along the dotted line at the edge of the unit cell. The electric field intensity above the grating remains relatively constant and low due to minimal transmission, whereas below the grating, it exhibits an interference pattern.

To gain a comprehensive understanding of the height-dependence of the absorbance, we simulated the dependence of the reflectance and absorbance spectra on the pillar height *d* using water as a model analyte inside the ITS device. Figure 2(d) illustrates the typical reflectance spectra for the devices with three different pillar heights *d* = 200 nm, 1000 nm and 2000 nm. The water absorption line at *ω*_H–O–H_is prominent in the reflectance spectra. The corresponding reflectance dip increases in intensity as the silica pillar height and consequently the sampled analyte thickness is increased. The dependence of the dip magnitude is plotted in Figure 2(e) as a function of *d*. The sin^2^(z) dependence of the field intensity resulting from the standing wave pattern gives rise to a nonlinear dependence of the absorbance intensity with the pillar height, as observed in figure 2(e).

Taken together, these findings enforce our interpretation of the reflectance spectrum of unpolarized light, *R*(*ω*) ≈ *R*_∥_/2 as inverse transflectance. In the rest of this work, we experimentally investigate how the inverse transflection absorbance spectra *A*(*ω*) = − log_10_(*R*_s_/*R*_bg_) (referred to as absorbance for brevity) can be used for sensing more complex analytes: proteins and live cells in aqueous environment. Here *R*_s_(*ω*) and *R*_bg_(*ω*) are the ITS measurements from the analyte-filled (sample) and water-filled (background) structures, respectively. The terms *reflectance* and *transflectance* will be used interchangeably in what follows because the former refers to the experimental technique while the latter – to the physical interpretation of the results.

## Results and discussions

The proposed device was fabricated through plasma-enhanced chemical vapor deposition (PECVD), electron beam lithography, electron beam evaporation of metals, and reactive ion etching (see the Experimental Section). The final fabricated samples used in the experiments described below comprise of thin gold gratings with a width of *w* = 200 nm atop of silica pillars with a height of *d* ≈ 830 nm, separated from each other by trenches with a width of *t* = *p* − *w*: see Figure 1(b) for an SEM image of a typical ITS device. Samples with three grating periodicities (*p* = 0.675 μm, 1 μm, and 1.35μm) were used.

### ITS of protein solutions

The ITS measurement of aqueous solutions of bovine serum albumin (BSA) of various concentrations were carried out, with the liquid samples completely filling the trench spaces, as well as the space above the gratings. Using unpolarized MIR light incident from the CaF_2_ side, we carried out the ITS measurements using a device with *p* = 1.0 μm. The transflectance spectra for the background (water only; blue line) and the 103 mg/mL BSA/water sample (red line) are plotted in Fig. 3(a), with the corresponding calculated absorbance shown in Fig. 3(b) as an orange line. The signature MIR spectral peaks of BSA^40–42^ are clearly visible, including the strongest amide I (*ω*_A−I_ ∼ 1,650 cm^−1^) and amide II (*ω*_A−II_ ∼ 1,548 cm^−1^) bands, as well as several weaker ones. Absorbance spectra for lower (42 mg/mL) and higher (164 mg/mL) BSA concentrations are shown in Figure 3(b) for comparison. The plot of the amide II peak integrated intensity shown in Figure 3(c) exhibits linear dependence on the BSA concentration in accordance with Beer Lambert Law. The results of our measurements using ITS devices filled with BSA water solutions demonstrate their utility for MIR spectroscopy of complex liquid analytes, with the measurement setup reminiscent of ATR-FTIR spectroscopy.

**Figure 3:**
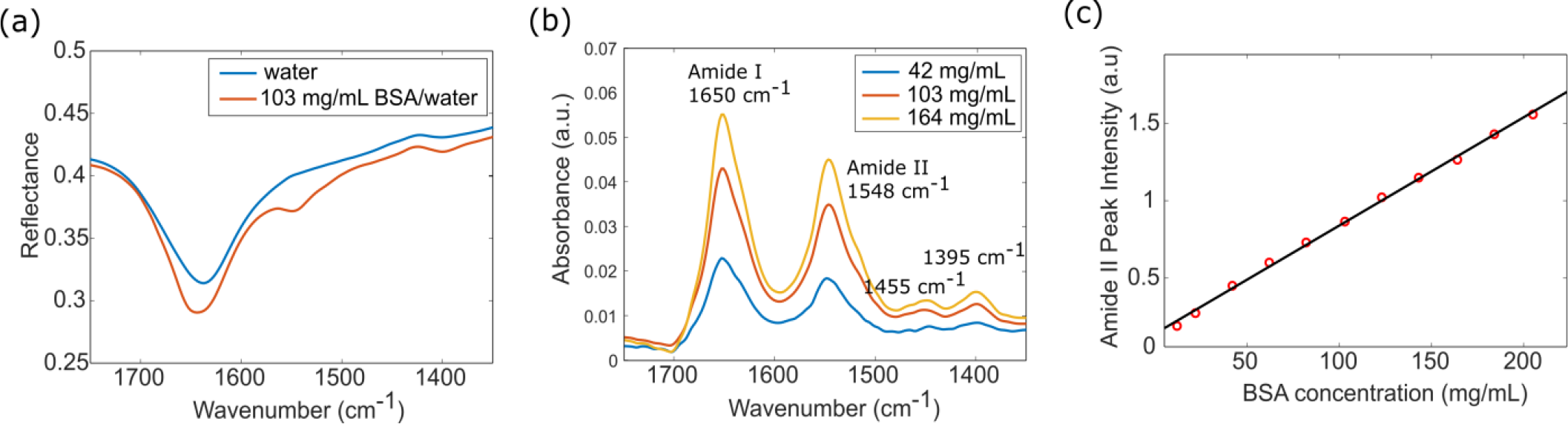
Experimentally measured bovine serum albumin (BSA) spectra using ITS. (a) Reflectance spectra measured from the nanograting with water and 103 mg/mL BSA solution. (b) Absorbance spectra of BSA in water with concentrations: 42 mg/mL, 103 mg/mL and 164 mg/mL showing several protein-related vibrational peaks. (c) Variation of the integrated peak intensity of the Amide II peak with BSA solution concentration. Nanograting parameters (see Fig.1(a) for geometry definitions): *p* = 0. 675 μm, *d* = 830 nm, and *w* = 200 nm.

### ITS of live cells: general considerations

Next, we demonstrate that ITS nanogratings can be applied to MIR spectroscopy of live cells. Compared with aqueous solution of proteins, IR spectra of cells are more complex to measure and interpret because of the heterogeneous distribution of different biomolecules within each cell, as well as their fairly complex (and frequently poorly understood) morphology. For example, it is not clear *a priori* whether a cell can be sufficiently deformed to penetrate into the trenches between the silica pillars. As Fig.2(b) illustrates, the very success of the nanograting-based ITS depends on such deformations because they ensure sufficient overlap with the MIR optical field inside the trenches.

Cellular interactions with nano-topographies have recently received considerable attention because of the potential effects of three-dimensional (3D) nano-structures on cell adhesion and morphology,^43,44^ penetration of cell membrane for cargo delivery,^45^ electroporation of the cellular membrane,^46,47^ as well as modulation of cell membrane curvatures.^48–51^ Interaction of adherent cells with micrometer- and nanometer-scale grooves and ridges – similar to the ones depicted in Fig.1, but without metallic strips on top – has been studied even earlier in the context of cell alignment and contact guidance.^52,53^ Crucially, it has been recently reported that with the appropriate width and periodicity of the nanostructures, many cell types can deform and tightly wrap around the vertical structures by closely following the nano-topography of the surface.^48–51^ Moreover, critical cell functions such as adhesion and membrane integrity, have been shown not to be impaired by the tight wrapping of the cellular membrane around 3D nanostructures.^54^ Nanostructured substrates have long been recognized as distinct cell culture dishes because of the effect of the topography on cellular attachment, division, and proliferation.^55^ High-aspect silicon-based nanostructures have also been used for enhancing the capture rate of rare cells owing to the local topographic interactions between the nanostructured substrates and nanoscale components of the cellular surface.^56,57^ The established fact of numerous cell types adhering closely to nanostructures by wrapping themselves closely around each 3D (elevated) grating structure lead us to surmise that the cells can completely take up the space inside the trenches, thus overlapping with the probing MIR light. Therefore, the effect of the cells lowering themselves into the trenches could be measured by the ITS device as a function of time.

To validate the effect of cell membrane deformation and cellular matter penetration into the nano-grooves, we first cultured A431 (human epidermoid carcinoma) cells on the 3D ITS nanostructure shown in Fig.1 and studied the membrane deformation using confocal florescence microscopy. Cells were stained with CellMask plasma membrane stain and imaged at different planes 1 and 2. The two imaged planes are shown in Figure 4(a) as dashed lines: plane 1 is near the bottom of the silica pillars while plane 2 is at the gold grating. Near the bottom of the silica pillars (see Figure 4(c): imaged plane 1 approximately 700 nm below the gold grating), the stained plasma membrane inside the trenches appears bright. Conversely, the silica pillars appear as periodic dark stripes because they naturally exclude the cellular material. This observation confirms that the cells are indeed curving around the silica pillars, reaching into the trenches. In imaged plane 2, located near the gold grating (see Figure 4(d)), we see bright green periodic stripes in the same locations as the dark stripes in Figure 4(c), corresponding to the location of the gold grating lines. This is clearly seen in Figure 4(b), where the pixel intensity values along the white dotted lines are plotted for these two planes. The brighter fluorescence signal at gold gratings is attributed to the metal-enhanced fluorescence effect, where fluorescence emission near metal surfaces is enhanced due to the coupling to localized surface plasmons and increased radiative decay rate of the fluorophores.^58^ Taken together, the fluorescence images shown in Figure 4 demonstrate that cellular material is indeed present above the nanograting, as well as deep inside the nanotrenches.

**Figure 4:**
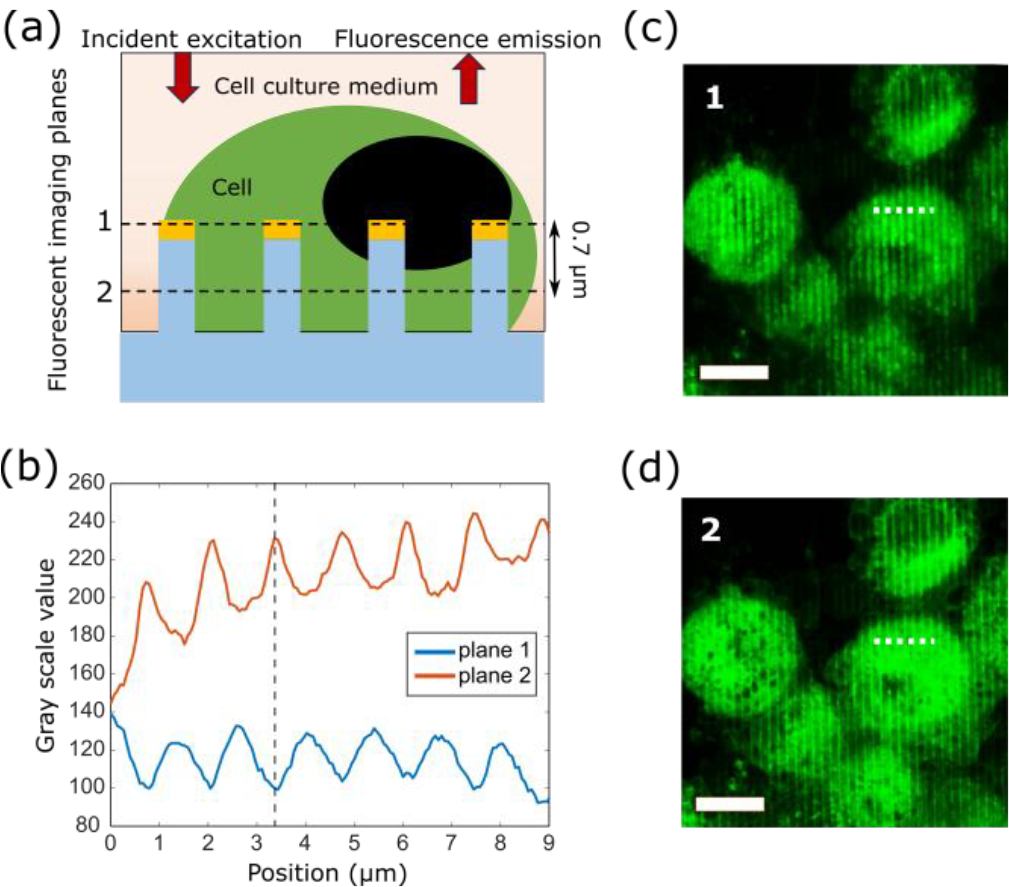
Confocal fluorescence microscopy image of A431 cells grown on the ITS device. Cells are stained with CellMask plasma membrane stain. (a) Device schematic, with 1 and 2 z-slice planes marked as dashed lines. (b) Fluorescence intensity along the white dashed lines marked in figure (c) and (d), showing that the intensity dips in plane 1 correspond to intensity peaks in plane 2. (c) and (d) Fluorescence images from planes 1 and 2, respectively. Dark stripes in (c): silica pillars locations (no cells). Bright stripes in (d): gold strips positions (metalenhanced fluorescence). Scale bar: 10 μm for all fluorescence images. ITS device parameters: *p* = 1. 35 μm, *d* = 830 nm, and *w* = 200 nm.

### ITS of live cells: endpoint measurements

Because the cells extending into the trenches can be interrogated using ITS, we proceeded to characterize the MIR spectra of live cells cultured atop of the nanogratings using FTIR spectroscopy: see Fig.5(a) for the measurement setup. Specifically, live cells were cultured inside microwells covered underneath by nanograting-containing CaF_2_ substrates. The cells were probed from the CaF_2_ (bottom) side while they were immersed in cell culture medium. The measured reflectance spectra, under normally incident unpolarized light, are shown in figure 5(b) for cells grown on nanogratings with 3 different periodicities: *p* = 0.675 μm, 1.0 μm & 1.35 μm. We observe that, except around the water absorption peak at *ω*_H–O–H_∼ 1,650 cm^−1^, the grating with *p* = 0.675 μm consistently provides reflectance in the 0.38 < *R*(*ω*) < 0.47 range across the entire measured wavenumber range 1,300 cm^−1^ < *ω* < 3,000 cm^−1^. However, the reflectance steadily decreases with increasing wavenumbers *ω* for the gratings with *p* = 1.0 μm or 1.35 μm. Consistently with Equation 1, this observation is attributed to increased transmission of the grating-polarized light at shorter wavelengths.

**Figure 5:**
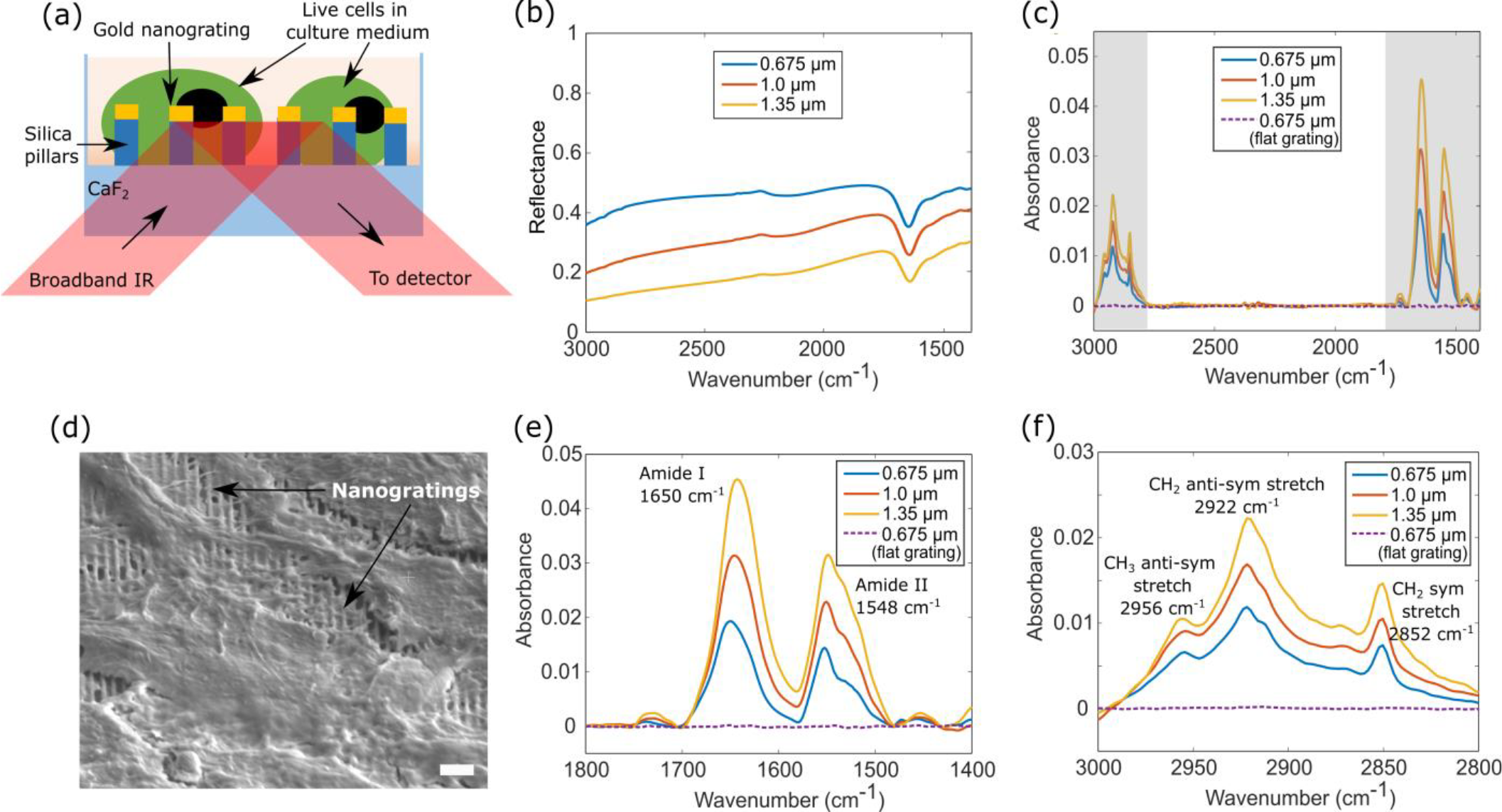
IR spectroscopy of live cells using the ITS device (a) Schematic of the measurement setup used for cell characterization. (b) Inverted transflectance and (c) the corresponding absorbance spectra of A431 cells grown on elevated gratings with 3 grating periodicities (blue: *p* = 0. 675 μm, red: *p* = 1. 0 μm, yellow: *p* = 1. 35 μm) and flat grating (dotted purple: *p* = 0. 675 μm). (d) SEM image of fixed and dried A431 cells grown on the nanograting (*p* = 0. 675 μm). Scale bar: 2 μm. (e) and (f): Enlarged absorbance spectra corresponding to the spectral regions grey-shaded in (c) attributed to (e) proteins (amide I and II) and (f) lipids (CH_2_/CH_3_), respectively.

On the other hand, increasing the periodicity while keeping the grating width *w* fixed increases the fractional volume of cells *v*_cell_∼ *t*/ *p* = 1 − *w*/ *p* in the trenches, resulting in their larger overlap with the optical field and stronger absorbance. Therefore, the optimal periodicity for broadband reflectance measurement requires a balance between maintaining sufficiently high reflectance (i.e. small *p*) and large *v*_cell_(i.e. large trench width *t*). For example, gratings with *p* > 1 μm are not suitable for measuring small changes, especially for the CH_2_/CH_3_ stretching modes attributed to lipids (*λ* ∼ 3.3μm) because the overall reflectance of the ITS device is strongly reduced (e.g., *R* ∼ 0.12 for *p* = 1.35μ*m*).

The close-ups for the regions of interest (proteins and lipids bands) of the cellular absorbance *A*_cell_(*ω*) plotted in Figures 5(e,f) provide a clear representation of the cellular vibrational spectra. They are very similar to the absorbance spectra measured in typical FTIR measurements of cells, including in the transmission and ATR-FTIR configuration, as well as our previous metasurface-based measurement using MEIRS.^1,22,27,59,60^ Here, the absorbance for the cellular measurements is defined as *A*_cell_(*ω*) = −log_10_(*R*(*ω*)^cell^/*R*(*ω*)^med^), where *R*(*ω*)^cell^ and *R*(*ω*)^med^ are the transflectance spectra measured from the nanograting with live cells in culture medium and from the cells-free culture medium, respectively. The dominant features observed in the cellular absorbance spectra include amide I (*ω*_A−I_ ∼ 1,650 cm^−1^) and amide II (*ω*_A−II_ ∼ 1,548 cm^−1^) peaks attributable to proteins, as well as the CH_2_ symmetric stretching 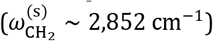, CH_2_ anti-symmetric stretching 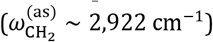 and CH_3_ antisymmetric stretching 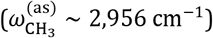 attributed to lipids. We note that while we have chosen to use the reflectance *R*(*ω*)^med^ from a medium-filled ITS device as a background signal, using water-filled structures as the background would also be acceptable because we did not detect any of the four characteristic frequencies (amides or CH stretches) in *R*(*ω*)^med^ (not shown).

### ITS of live cells: Real-time measurements of cell adhesion

To demonstrate the utility of the ITS device for long-term kinetic measurement of live cells, we spectroscopically characterize cellular adhesion as a function of time using the nanograting with periodicity *p* = 0.675 μm. The reflectance spectra were measured every 2 minutes for 5 hours as the cells were seeded and adhered to the nanograting structures. To observe the evolution in protein and lipid absorbance signals over time, we analyzed the integrated peak intensity of the Amide II peak at *ω*_A−II_ and the CH_2_ symmetric stretching mode at 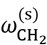, respectively. These intensities were plotted as a function of time in figures 6(c) and (d), respectively. Our analysis reveals similar trends in the temporal signal evolution for both peaks. Specifically, the signals grow rapidly during the first 140 minutes, and then their growth rate slows down – consistent with our previous observations of cell adhesion using a plasmonic metasurface.^29^

**Figure 6:**
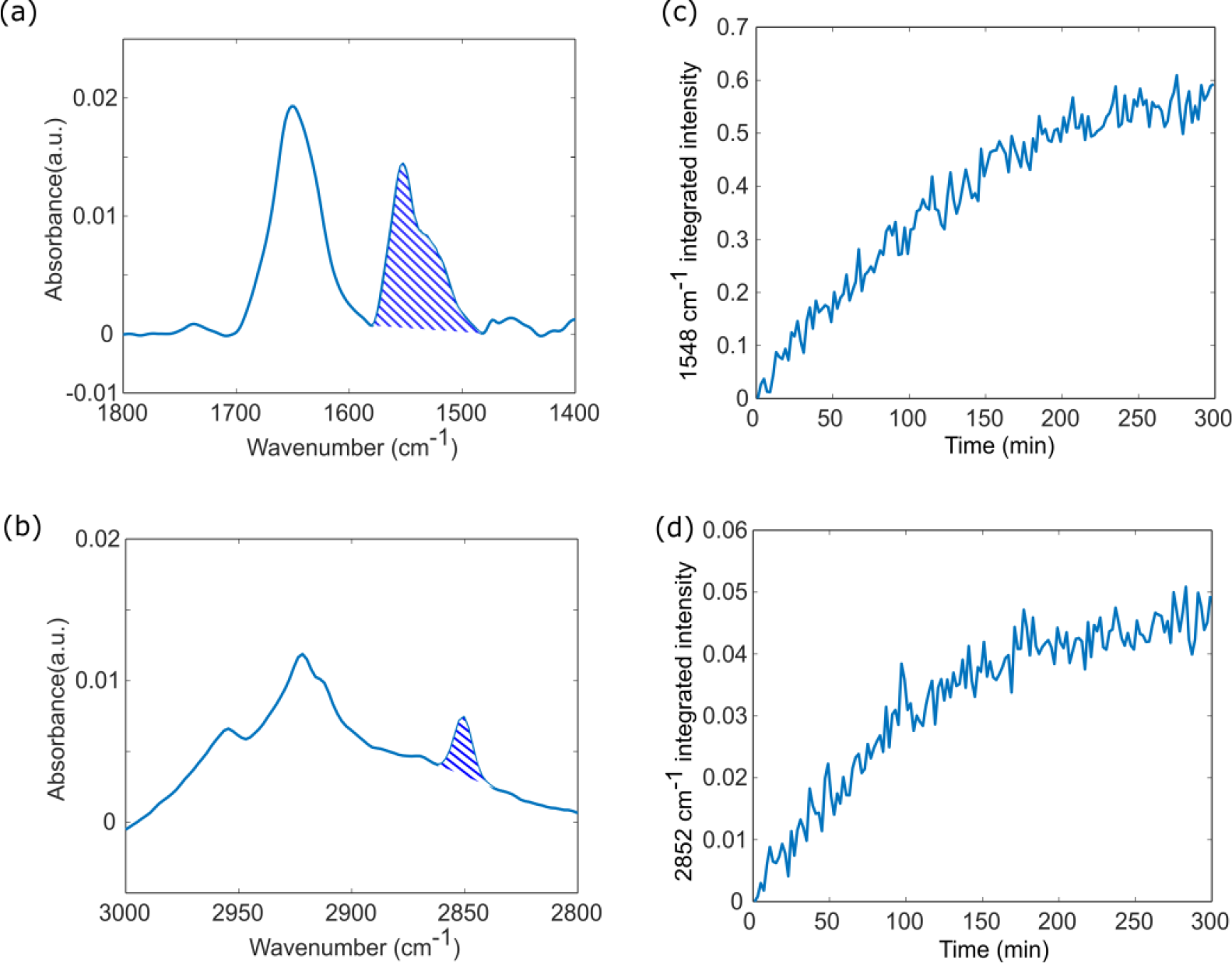
Time evolution of live A431 cell adhesion on the ITS device. (a) and (b) Absorbance spectra corresponding to protein and lipid bands respectively, measured using grating with periodicity *p* = 0. 675 μm, pillar height *d* = 830 nm and grating width *w* = 200 nm. Shaded regions indicate the region over which the peaks were integrated (c) Time evolution of the integrated peak intensity of amide II band, representing protein signal evolution with cellular adhesion. (d) Time evolution of the integrated peak intensity of CH_2_ symmetric stretching, representing lipid signal evolution with cellular adhesion.

Future studies will be extended to investigating whether different surface coatings affect cellular adhesion and growth rate on such nanocontoured substrates. Typically, cell adhesion on flat surfaces is heavily reliant on the presence of extra-cellular matrix (ECM) adhesion proteins, such as collagen and fibronectin, for integrin mediated cell adhesion. However, on nanostructured surfaces, it has been reported that clathrin endocytosis complexes forming around the surface protrusions play an important role in the cellular interaction with nanostructured surfaces. Specifically, clathrin-mediated endocytosis has been implicated in the formation of membrane curvature around nanostructures.^61^ By examining the effects of various surface coatings on cellular adhesion and proliferation rates, we can gain insights into how surface topography influences cell behavior (e.g., by modulating cell membrane curvatures^48–51^), and potentially develop strategies for enhancing or inhibiting cell adhesion and growth on nanocontoured substrates.

## Conclusion

In this work, we introduced a novel MIR spectroscopy-based live cell biosensing platform – Inverted Transflectance Spectroscopy using elevated nanogratings. We show that ITS devices produce broadband signals and enable MIR spectroscopy in the 1400 cm^-1^ -3000 cm^-1^ spectral range. ITS can be used to characterize analytes in solution or liquid form as we demonstrate through the IR spectroscopy of BSA solutions. Further, adherent cells can be measured as well, taking advantage of the fact that the cells can deform around the elevated nanogratings and extend into the trenches below. We demonstrated this by characterizing the adhesion kinetics of live A431 cells through MIR spectroscopy.

Currently, the spectral range of the presented ITS device is limited by the absorption caused by the silica layer. To extend the useful spectral range below 1400 cm^-1^, alternative materials such as Al_2_O_3_, which are more transparent in the relevant spectral region, could be used to fabricate the dielectric pillars.

The nano-grating based device described here is somewhat similar to metasurface-based device reported by our group previously,^28,29,62^ but these two devices have different sensing volumes and can be complementary to each other. Since the metasurface sensing volume is limited by the near-field decay of the plasmonic hotspots (roughly 100 nm), it is limited to the sensing of cellular membrane and cellular structures around the membrane. On the other hand, the ITS device described in this work is based on transflectance measurement. As long as the cells can extend into the trenches below the nano-grating, they can be probed much beyond the cell membrane. Cells can also undergo significant deformation of their nucleus around such vertical nanostructures.^46,63^ Such nuclear deformation would enable MIR spectroscopy of the nucleus: an important measurement that will be carried out in our future work. For non-adherent cells the ITS device can in turn be used to monitor secretion of cell metabolites and uptake of nutrients.

## Experimental Section

### Device Fabrication

A 12.5 mm x 12.5 mm x 0.5 mm calcium fluoride substrate was coated with a 1 μm thick silica thin film using plasma-enhanced chemical vapor deposition (PECVD). Polymethyl methacrylate (PMMA) e-beam resist was spin-coated on to the surface with the silica thin film and nanograting pattern were defined through e-beam lithography. Next, a 5 nm Cr adhesion layer, 70 nm Au layer and 18 nm of Cr mask layer was evaporated onto the resulting structure. The grating pattern was transferred on to the Au layer through lift-off in an acetone bath. Reactive Ion Etch (RIE) -with a mixture of CF_4_ and Ar, was used to etch down the silica using the chromium layer as a hard mask. The silica layer was etched down to form the elevated grating structure of height 830 nm, with about 170 nm of unetched silica left at the bottom. The fabricated gratings had a width of 200 nm and periodicities of 0.675 μm, 1 μm and 1.35 μm.

### Numerical simulation

The reflectance spectrum and near field profile of the nanograting sample are simulated using COMSOL Multiphysics, a commercially available software using finite elements method. The geometric parameters are taken from sample SEM images, with gold thickness of 70 nm. The grating is simulated in a unit cell with Floquet boundary conditions in the x and y directions. The grating structure is simulated with water on the top and supported by CaF_2_ on the bottom, both layers with thickness equal to at least one wavelength. The wave excitation is simulated using a periodic port, where plane waves are normally incident from the CaF_2_ side. To determine the 1648 cm^-1^ peak absorbance in Figure 2(e), we integrate the area under the simulated reflectance dips as depicted in Figure 2(d).

### Sample preparation for cell measurements

Before seeding the cells for incubation or adhesion measurement, the nanograting device was sterilized with 70% ethanol, incubated overnight in 10 μg/mL fibronectin solution in Dulbecco’s phosphate-buffered saline (DPBS) and washed with DPBS twice. Human epidermoid carcinoma cell line A431 cells were obtained from the American Type Culture Collection (ATCC) and cultured in Dulbecco’s Modified Eagle Medium (DMEM) supplemented with 10% fetal bovine serum (FBS) and 1% penicillin-streptomycin. For cellular endpoint measurements, A431 cells were detached from culture flask using 0.25% trypsin-EDTA and seeded on the fibronectin-coated nanograting device. The cells were incubated on the nanograting device in DMEM + 10% FBS until confluent coverage (>95%) was achieved. Prior to measurement, the culture medium was exchanged to serum-free Leibovitz’s L15 medium for buffering in ambient air condition.

For the cell adhesion experiment, A431 cells were grown in culture flasks in an incubator and the nanograting device was prepared by coating it with fibronectin. The cells were removed from the culture flask by trypsinization and then seeded on the nanograting device in serum-free Leibovitz’s L15 medium.

### FTIR spectroscopy

FTIR spectroscopy was performed using a Bruker Vertex v70 FTIR spectrometer. All measurements were performed at room temperature in ambient air conditions, using unpolarized light.

**Cellular endpoint measurement and measurement of BSA concentration** was made using Bruker Hyperion 3000 infrared microscope, fitted with a reflective Cassegrain objective (15X, NA = 0.4) and a mercury-cadmium-telluride (MCT) detector. For the endpoint measurements the absorbance was calculated from averaging 2 measurements on samples with the same parameters.

**Cellular adhesion measurement** was done using an in-house inverted mid-infrared microscope which was built as an external add-on to the FTIR spectrometer and is described in detail elsewhere^29^. Measurements were made in reflectance mode, through the CaF2 substrate. FTIR spectra was collected at the rate of 1 acquisition every 2 minutes. The time series was calculated by averaging data from 3 measurements on samples with the same parameters. To correct for different concentrations of cells covering the nanograting pixel, the absorbance intensities were scaled such that the intensities at *t* = 300 mins match.

For all the spectroscopic measurements, background spectra are collected from a patch of gold fabricated on the same chip as the nanogratings and the FTIR spectra were collected with 100 averaging for both background and sample, at 4 cm^-1^ resolution. As post-processing, water vapor spectra were subtracted from the acquired spectra using Bruker OPUS software. Baseline correction was used to remove baseline offsets in the calculated absorbance.

### Fluorescence staining and Confocal Fluorescence microscopy

Fresh working solution (5 μg /mL) of CellMask plasma membrane stain was prepared by mixing the concentrated solution (5 mg/mL) in PBS. A431 cells grown on the nanograting device were submerged in the working solution for 10 mins at 37 °C. The staining solution is removed from the sample at the sample is washed 3 times in PBS. Finally, the cells are fixed with 4% formaldehyde for 15 mins at room temperature. For fluorescence imaging we use Zeiss LSM710 confocal microscope. The microscope imaging setup had a focal depth of 0.7 μm.

## Author Contributions

The manuscript was written through contributions of all authors. All authors have given approval to the final version of the manuscript.

## ACKNOWLEDGMENT

The research reported here was supported by the National Cancer Institute of the National Institutes of Health under award number R21CA251052 and by the National Institute of General Medical Sciences of the National Institutes of Health under award number R21GM138947. This work was performed in part at the Cornell NanoScale Science and Technology Facility (CNF), a member of the National Nanotechnology Coordinated Infrastructure (NNCI), which is supported by the National Science Foundation (Grant NNCI-2025233). This work was also performed in part at the BRC Imaging Facilty at Cornell, which is supported by the National Institute of Health (Grant NIH S10RR025502). The authors acknowledge the use of facilities and instrumentation supported by NSF through the Cornell University Materials Research Science and Engineering Center DMR-1719875.

## REFERENCES

(1) Baker, M. J.; Trevisan, J.; Bassan, P.; Bhargava, R.; Butler, H. J.; Dorling, K. M.; Fielden, P. R.; Fogarty, S. W.; Fullwood, N. J.; Heys, K. A.; Hughes, C.; Lasch, P.; Martin-Hirsch, P. L.; Obinaju, B.; Sockalingum, G. D.; Sulé-Suso, J.; Strong, R. J.; Walsh, M. J.; Wood, B. R.; Gardner, P.; Martin, F. L. Using Fourier Transform IR Spectroscopy to Analyze Biological Materials. Nat Protoc 2014, 9 (8), 1771–1791. 10.1038/nprot.2014.110.

(2) Jackson, M.; Mantsch, H. H. The Use and Misuse of FTIR Spectroscopy in the Determination of Protein Structure. Crit Rev Biochem Mol Biol 1995, 30 (2), 95–120. 10.3109/10409239509085140.

(3) Yang, H.; Yang, S.; Kong, J.; Dong, A.; Yu, S. Obtaining Information about Protein Secondary Structures in Aqueous Solution Using Fourier Transform IR Spectroscopy. Nat Protoc 2015, 10 (3), 382–396. 10.1038/nprot.2015.024.

(4) Wu, C.; Khanikaev, A. B.; Adato, R.; Arju, N.; Yanik, A. A.; Altug, H.; Shvets, G. Fano-Resonant Asymmetric Metamaterials for Ultrasensitive Spectroscopy and Identification of Molecular Monolayers. Nature Materials 2011 11:1 2011, 11 (1), 69–75. 10.1038/NMAT3161.

(5) Delbeck, S.; Heise, H. M. FT-IR versus EC-QCL Spectroscopy for Biopharmaceutical Quality Assessment with Focus on Insulin—Total Protein Assay and Secondary Structure Analysis Using Attenuated Total Reflection. Anal Bioanal Chem 2020, 412 (19), 4647. 10.1007/S00216-020-02718-1.

(6) Sala, A.; Spalding, K. E.; Ashton, K. M.; Board, R.; Butler, H. J.; Dawson, T. P.; Harris, D. A.; Hughes, C. S.; Jenkins, C. A.; Jenkinson, M. D.; Palmer, D. S.; Smith, B. R.; Thornton, C. A.; Baker, M. J. Rapid Analysis of Disease State in Liquid Human Serum Combining Infrared Spectroscopy and “Digital Drying.” J Biophotonics 2020, 13 (9), e202000118. 10.1002/JBIO.202000118.

(7) Flower, K. R.; Khalifa, I.; Bassan, P.; Démoulin, D.; Jackson, E.; Lockyer, N. P.; McGown, A. T.; Miles, P.; Vaccari, L.; Gardner, P. Synchrotron FTIR Analysis of Drug Treated Ovarian A2780 Cells: An Ability to Differentiate Cell Response to Different Drugs? Analyst 2011, 136 (3), 498–507. 10.1039/C0AN00564A.

(8) Gough, K. M.; Tzadu, L.; Kastyak, M. Z.; Kuzyk, A. C.; Julian, R. L. Theoretical and Experimental Considerations for Interpretation of Amide I Bands in Tissue. Vib Spectrosc 2010, 53 (1), 71–76. 10.1016/J.VIBSPEC.2010.01.015.

(9) Gazi, E.; Dwyer, J.; Lockyer, N. P.; Miyan, J.; Gardner, P.; Hart, C. A.; Brown, M. D.; Clarke, N. W. A Study of Cytokinetic and Motile Prostate Cancer Cells Using Synchrotron-Based FTIR Microspectroscopic Imaging. Vib Spectrosc 2005, 38 (1–2), 193–201. 10.1016/J.VIBSPEC.2005.02.026.

(10) Alcaráz, M. R.; Schwaighofer, A.; Kristament, C.; Ramer, G.; Brandstetter, M.; Goicoechea, H.; Lendl, B. External-Cavity Quantum Cascade Laser Spectroscopy for Mid-IR Transmission Measurements of Proteins in Aqueous Solution. Anal Chem 2015, 87 (13), 6980–6987. 10.1021/ACS.ANALCHEM.5B01738/SUPPL_FILE/AC5B01738_SI_001.PDF.

(11) Akhgar, C. K.; Ramer, G.; Żbik, M.; Trajnerowicz, A.; Pawluczyk, J.; Schwaighofer, A.; Lendl, B. The Next Generation of IR Spectroscopy: EC-QCL-Based Mid-IR Transmission Spectroscopy of Proteins with Balanced Detection. Anal Chem 2020, 92 (14), 9901–9907. 10.1021/acs.analchem.0c01406.

(12) Chon, B.; Xu, S.; Lee, Y. J. Compensation of Strong Water Absorption in Infrared Spectroscopy Reveals the Secondary Structure of Proteins in Dilute Solutions. Anal Chem 2021, 93 (4), 2215–2225. 10.1021/acs.analchem.0c04091.

(13) Kazarian, S.; Chan, S. K. L. A.; Kazarian, S. G.; Chan, K. L. A. Attenuated Total Reflection Fourier-Transform Infrared (ATR-FTIR) Imaging of Tissues and Live Cells. Chem Soc Rev 2016, 45 (7), 1850–1864. 10.1039/C5CS00515A.

(14) Spalding, K.; Bonnier, F.; Bruno, C.; Blasco, H.; Board, R.; Benz-de Bretagne, I.; Byrne, H. J.; Butler, H. J.; Chourpa, I.; Radhakrishnan, P.; Baker, M. J. Enabling Quantification of Protein Concentration in Human Serum Biopsies Using Attenuated Total Reflectance – Fourier Transform Infrared (ATR-FTIR) Spectroscopy. Vib Spectrosc 2018, 99, 50–58. 10.1016/J.VIBSPEC.2018.08.019.

(15) Yamaguchi, R. T.; Hirano-Iwata, A.; Kimura, Y.; Niwano, M.; Miyamoto, K. I.; Isoda, H.; Miyazaki, H. In Situ Real-Time Monitoring of Apoptosis on Leukemia Cells by Surface Infrared Spectroscopy. J Appl Phys 2009, 105 (2). 10.1063/1.3068203/284268.

(16) Mordechai, S.; Sahu, R. K.; Hammody, Z.; Mark, S.; Kantarovich, K.; Guterman, H.; Podshyvalov, A.; Goldstein, J.; Argov, S. Possible Common Biomarkers from FTIR Microspectroscopy of Cervical Cancer and Melanoma. J Microsc 2004, 215 (Pt 1), 86–91. 10.1111/J.0022-2720.2004.01356.X.

(17) Argov, S.; Ramesh, J.; Salman, A.; Sinelnikov, I.; Goldstein, J.; Guterman, H.; Mordechai, S. Diagnostic Potential of FTIR Microspectroscopy and Advanced Computational Methods in Colon Cancer Patients. 10.1117/1.1463051 2002, 7 (2), 248–254. 10.1117/1.1463051.

(18) Diem, M.; Miljković, M.; Bird, B.; Chernenko, T.; Schubert, J.; Marcsisin, E.; Mazur, A.; Kingston, E.; Zuser, E.; Papamarkakis, K.; Laver, N. Applications of Infrared and Raman Microspectroscopy of Cells and Tissue in Medical Diagnostics: Present Status and Future Promises. Spectroscopy (New York) 2012, 27 (5–6), 463–496. 10.1155/2012/848360.

(19) Derenne, A.; Gasper, R.; Goormaghtigh, E. The FTIR Spectrum of Prostate Cancer Cells Allows the Classification of Anticancer Drugs According to Their Mode of Action. Analyst 2011, 136 (6), 1134–1141. 10.1039/C0AN00872A.

(20) Chan, K. L. A.; Fale, P. L. V. Label-Free in Situ Quantification of Drug in Living Cells at Micromolar Levels Using Infrared Spectroscopy. Anal Chem 2014, 86 (23), 11673–11679. 10.1021/AC503915C/SUPPL_FILE/AC503915C_SI_001.PDF.

(21) Fale, P. L.; Altharawi, A.; Chan, K. L. A. In Situ Fourier Transform Infrared Analysis of Live Cells’ Response to Doxorubicin. Biochimica et Biophysica Acta (BBA) - Molecular Cell Research 2015, 1853 (10), 2640–2648. 10.1016/J.BBAMCR.2015.07.018.

(22) Altharawi, A.; Rahman, K. M.; Chan, K. L. A. Towards Identifying the Mode of Action of Drugs Using Live-Cell FTIR Spectroscopy. Analyst 2019, 144 (8), 2725–2735. 10.1039/C8AN02218F.

(23) Aonuma, Y.; Kondo, Y.; Hirano-Iwata, A.; Nishikawa, A.; Shinohara, Y.; Iwata, H.; Kimura, Y.; Niwano, M. Label-Free and Real Time Monitoring of Adipocyte Differentiation by Surface Infrared Spectroscopy. Sens Actuators B Chem 2013, 176, 1176–1182. 10.1016/J.SNB.2012.10.030.

(24) Grenci, G.; Birarda, G.; Mitri, E.; Businaro, L.; Pacor, S.; Vaccari, L.; Tormen, M. Optimization of Microfluidic Systems for IRMS Long Term Measurement of Living Cells. Microelectron Eng 2012, 98, 698–702. 10.1016/J.MEE.2012.05.049.

(25) Vaccari, L.; Birarda, G.; Businaro, L.; Pacor, S.; Grenci, G. Infrared Microspectroscopy of Live Cells in Microfluidic Devices (MD-IRMS): Toward a Powerful Label-Free Cell-Based Assay. Anal Chem 2012, 84 (11), 4768–4775. 10.1021/AC300313X/SUPPL_FILE/AC300313X_SI_001.PDF.

(26) Kelp, G.; Arju, N.; Lee, A.; Esquivel, E.; Delgado, R.; Yu, Y.; Dutta-Gupta, S.; Sokolov, K.; Shvets, G. Application of Metasurface-Enhanced Infra-Red Spectroscopy to Distinguish between Normal and Cancerous Cell Types. Analyst 2019, 144 (4), 1115–1127. 10.1039/C8AN01433G.

(27) Kelp, G.; Li, J.; Lu, J.; DiNapoli, N.; Delgado, R.; Liu, C.; Fan, D.; Dutta-Gupta, S.; Shvets, G. Infrared Spectroscopy of Live Cells from a Flowing Solution Using Electrically-Biased Plasmonic Metasurfaces. Lab Chip 2020, 20 (12), 2136–2153. 10.1039/C9LC01054H.

(28) Huang, S. H.; Li, J.; Fan, Z.; Delgado, R.; Shvets, G. Monitoring the Effects of Chemical Stimuli on Live Cells with Metasur-face-Enhanced Infrared Reflection Spectroscopy. Lab Chip 2021, 21 (20), 3991–4004. 10.1039/D1LC00580D.

(29) Huang, S. H.; Sartorello, G.; Shen, P. T.; Xu, C.; Elemento, O.; Shvets, G. Metasurface-Enhanced Infrared Spectroscopy in Multiwell Format for Real-Time Assaying of Live Cells. Lab Chip 2023. 10.1039/D3LC00017F.

(30) Shen, P. T.; Huang, S. H.; Huang, Z.; Wilson, J. J.; Shvets, G. Probing the Drug Dynamics of Chemotherapeutics Using Metasur-face-Enhanced Infrared Reflection Spectroscopy of Live Cells. Cells 2022, 11 (10), 1600. 10.3390/CELLS11101600/S1.

(31) Rodrigo, D.; Tittl, A.; Ait-Bouziad, N.; John-Herpin, A.; Limaj, O.; Kelly, C.; Yoo, D.; Wittenberg, N. J.; Oh, S. H.; Lashuel, H. A.; Altug, H. Resolving Molecule-Specific Information in Dynamic Lipid Membrane Processes with Multi-Resonant Infrared Metasurfaces. Nature Communications 2018 9:1 2018, 9 (1), 1–9. 10.1038/S41467-018-04594-X.

(32) H. Lamb. On the Reflection and Transmission of Electric Waves by a Metallic Grating. Proc. London Math. Soc. 1898, 29 (523).

(33) Kitamura, R.; Pilon, L.; Jonasz, M. Optical Constants of Silica Glass from Extreme Ultraviolet to Far Infrared at near room Temperature. Applied Optics, Vol. 46, Issue 33, pp. 8118–8133 2007, 46 (33), 8118–8133. 10.1364/AO.46.008118.

(34) Kotanen, C. N.; Gabriel Moussy, F.; Carrara, S.; Guiseppi-Elie, A. Infrared Spectroscopy of Membrane Lipids. Encyclopedia of Biophysics 2013, 1074–1081. 10.1007/978-3-642-16712-6_558.

(35) Barth, A. Infrared Spectroscopy of Proteins. Biochimica et Biophysica Acta (BBA) - Bioenergetics 2007, 1767 (9), 1073–1101. 10.1016/J.BBABIO.2007.06.004.

(36) Poonprasartporn, A.; Chan, K. L. A. Live-Cell ATR-FTIR Spectroscopy as a Novel Bioanalytical Tool for Cell Glucose Metabolism Research. Biochimica et Biophysica Acta (BBA) - Molecular Cell Research 2021, 1868 (7), 119024. 10.1016/J.BBAMCR.2021.119024.

(37) Brooke, H.; Bronk, B. V.; McCutcheon, J. N.; Morgan, S. L.; Myrick, M. L. A Study of Electric Field Standing Waves on Reflection Microspectroscopy of Polystyrene Particles. 10.1366/000370209789806902 2009, 63 (11), 1293–1302. 10.1366/000370209789806902.

(38) Bassan, P.; Byrne, H. J.; Lee, J.; Bonnier, F.; Clarke, C.; Dumas, P.; Gazi, E.; Brown, M. D.; Clarke, N. W.; Gardner, P. Reflection Contributions to the Dispersion Artefact in FTIR Spectra of Single Biological Cells. Analyst 2009, 134 (6), 1171–1175. 10.1039/B821349F.

(39) Filik, J.; Frogley, M. D.; Pijanka, J. K.; Wehbe, K.; Cinque, G. Electric Field Standing Wave Artefacts in FTIR Micro-Spectroscopy of Biological Materials. Analyst 2012, 137 (4), 853–861. 10.1039/C2AN15995C.

(40) Schwaighofer, A.; Akhgar, C. K.; Lendl, B. Broadband La-ser-Based Mid-IR Spectroscopy for Analysis of Proteins and Monitoring of Enzyme Activity. Spectrochim Acta A Mol Biomol Spectrosc 2021, 253, 119563. 10.1016/J.SAA.2021.119563.

(41) Grdadolnik, J.; Chal, Y. M. Bovine Serum Albumin Observed by Infrared Spectrometry. II. Hydration Mechanisms and Interaction Configurations of Embedded H 2 O Molecules. 2000. 10.1002/1097-0282.

(42) Mittal, V.; Nedeljkovic, M.; Carpenter, L. G.; Khokhar, A. Z.; Chong, H. M. H.; Mashanovich, G. Z.; Bartlett, P. N.; Wilkinson, J. S. Waveguide Absorption Spectroscopy of Bovine Serum Albumin in the Mid-Infrared Fingerprint Region. ACS Sens 2019, 4 (7), 1749–1753. 10.1021/ACSSENSORS.9B00215/ASSET/IMAGES/LARGE/SE-2019-00215V_0004.JPEG.

(43) Kim, D. J.; Seol, J. K.; Lee, G.; Kim, G. S.; Lee, S. K. Cell Adhesion and Migration on Nanopatterned Substrates and Their Effects on Cell-Capture Yield. Nanotechnology 2012, 23 (39), 395102. 10.1088/0957-4484/23/39/395102.

(44) Zeng, Y.; Wong, S. T.; Teo, S. K.; Leong, K. W.; Chiam, K. H.; Yim, E. K. F. Human Mesenchymal Stem Cell Basal Membrane Bending on Gratings Is Dependent on Both Grating Width and Curvature. Scientific Reports 2018 8:1 2018, 8 (1), 1–13. 10.1038/s41598-018-24123-6.

(45) Peer, E.; Artzy-Schnirman, A.; Gepstein, L.; Sivan, U. Hollow Nanoneedle Array and Its Utilization for Repeated Administration of Biomolecules to the Same Cells. ACS Nano 2012, 6 (6), 4940–4946. 10.1021/NN300443H/SUPPL_FILE/NN300443H_SI_003.PDF.

(46) Caprettini, V.; Huang, J. A.; Moia, F.; Jacassi, A.; Gonano, C. A.; Maccaferri, N.; Capozza, R.; Dipalo, M.; De Angelis, F. Enhanced Raman Investigation of Cell Membrane and Intracellular Compounds by 3D Plasmonic Nanoelectrode Arrays. Advanced Science 2018, 5 (12). 10.1002/ADVS.201800560.

(47) Xie, C.; Lin, Z.; Hanson, L.; Cui, Y.; Cui, B. Intracellular Recording of Action Potentials by Nanopillar Electroporation. Nature Nanotechnology 2012 7:3 2012, 7 (3), 185–190. 10.1038/nnano.2012.8.

(48) Zhao, W.; Hanson, L.; Lou, H. Y.; Akamatsu, M.; Chowdary, P. D.; Santoro, F.; Marks, J. R.; Grassart, A.; Drubin, D. G.; Cui, Y.; Cui, B. Nanoscale Manipulation of Membrane Curvature for Probing Endocytosis in Live Cells. Nature Nanotechnology 2017 12:8 2017, 12 (8), 750–756. 10.1038/nnano.2017.98.

(49) Li, X.; Matino, L.; Zhang, W.; Klausen, L.; McGuire, A. F.; Lubrano, C.; Zhao, W.; Santoro, F.; Cui, B. A Nanostructure Platform for Live-Cell Manipulation of Membrane Curvature. Nature Protocols 2019 14:6 2019, 14 (6), 1772–1802. 10.1038/s41596-019-0161-7.

(50) Li, X.; Zhang, W.; Tsai, C. T.; Cui, B. Vertical Nanostructures for Probing Live Cells. Micro and Nano Systems for Biophysical Studies of Cells and Small Organisms 2021, 43–70. 10.1016/B978-0-12-823990-2.00003-9.

(51) Roy, A. R.; Zhang, W.; Jahed, Z.; Tsai, C. T.; Cui, B.; Moerner, W. E. Exploring Cell Surface-Nanopillar Interactions with 3D Super-Resolution Microscopy. ACS Nano 2022, 16 (1), 192–210. 10.1021/ACSNANO.1C05313/ASSET/IMAGES/LARGE/NN1C05313_0009.JPEG.

(52) Oakley, C.; Jaeger, N. A. F.; Brunette, D. M. Sensitivity of Fibroblasts and Their Cytoskeletons to Substratum Topographies: Topographic Guidance and Topographic Compensation by Micromachined Grooves of Different Dimensions; 1997; Vol. 234.

(53) Teixeira, A. I.; Abrams, G. A.; Bertics, P. J.; Murphy, C. J.; Nealey, P. F. Epithelial Contact Guidance on Well-Defined Micro- and Nanostructured Substrates. J Cell Sci 2003, 116 (10), 1881–1892. 10.1242/jcs.00383.

(54) Berthing, T.; Bonde, S.; Sørensen, C. B.; Utko, P.; Martinez, K. L.; Nygård, J. Intact Mammalian Cell Function on Semiconductor Nanowire Arrays: New Perspectives for Cell-Based Biosensing. Small 2011, 7 (5), 640–647. 10.1002/smll.201001642.

(55) Nomura, S.; Kojima, H.; Ohyabu, Y.; Kuwabara, K.; Miyauchi, A.; Uemura, T. Nanopillar Sheets as a New Type of Cell Culture Dish: Detailed Study of HeLa Cells Cultured on Nanopillar Sheets. Journal of Artificial Organs 2006, 9 (2), 90–96. 10.1007/s10047-006-0329-0.

(56) Liu, W. F.; Chen, C. S. Cellular and Multicellular Form and Function. Advanced Drug Delivery Reviews. November 10, 2007, pp 1319–1328. 10.1016/j.addr.2007.08.011.

(57) Wang, S.; Wang, H.; Jiao, J.; Chen, K. J.; Owens, G. E.; Kamei, K. I.; Sun, J.; Sherman, D. J.; Behrenbruch, C. P.; Wu, H.; Tseng, H. R. Three-Dimensional Nanostructured Substrates toward Efficient Capture of Circulating Tumor Cells. Angewandte Chemie - International Edition 2009, 48 (47), 8970–8973. 10.1002/anie.200901668.

(58) Jeong, Y.; Kook, Y. M.; Lee, K.; Koh, W. G. Metal Enhanced Fluorescence (MEF) for Biosensors: General Approaches and a Review of Recent Developments. Biosens Bioelectron 2018, 111, 102–116. 10.1016/J.BIOS.2018.04.007.

(59) Martin, F. L.; Kelly, J. G.; Llabjani, V.; Martin-Hirsch, P. L.; Patel, I. I.; Trevisan, J.; Fullwood, N. J.; Walsh, M. J. Distinguishing Cell Types or Populations Based on the Computational Analysis of Their Infrared Spectra. Nature Protocols 2010 5:11 2010, 5 (11), 1748–1760. 10.1038/nprot.2010.133.

(60) Mittal, S.; Yeh, K.; Suzanne Leslie, L.; Kenkel, S.; Kajdacsy-Balla, A.; Bhargava, R. Simultaneous Cancer and Tumor Microenvironment Subtyping Using Confocal Infrared Microscopy for All-Digital Molecular Histopathology. Proc Natl Acad Sci U S A 2018, 115 (25), E5651–E5660. 10.1073/PNAS.1719551115/SUPPL_FILE/PNAS.1719551115.SAPP.PDF.

(61) Cooper, G. M. Endocytosis. 2000.

(62) Kelp, G.; Kelp, G.; Kelp, G.; Li, J.; Lu, J.; Dinapoli, N.; Delgado, R.; Liu, C.; Fan, D.; Dutta-Gupta, S.; Dutta-Gupta, S.; Shvets, G. Infrared Spectroscopy of Live Cells from a Flowing Solution Using Electrically-Biased Plasmonic Metasurfaces. Lab Chip 2020, 20 (12), 2136–2153. 10.1039/C9LC01054H.

(63) Hanson, L.; Zhao, W.; Lou, H. Y.; Lin, Z. C.; Lee, S. W.; Chowdary, P.; Cui, Y.; Cui, B. Vertical Nanopillars for in Situ Probing of Nuclear Mechanics in Adherent Cells. Nature Nanotechnology 2015 10:6 2015, 10 (6), 554–562. 10.1038/nnano.2015.88.

